# Innate CD8αα^+^ cells and osteopontin promote ILC1-like intraepithelial lymphocyte homeostasis and intestinal inflammation

**DOI:** 10.1101/606616

**Authors:** Ali Nazmi, Kristen Hoek, Michael J. Greer, M. Blanca Piazuelo, Nagahiro Minato, Danyvid Olivares-Villagómez

**Author notes:** Corresponding author: Danyvid Olivares-Villagómez.; e-mail.

## Abstract

Innate CD8αα^+^ cells, also referred to as iCD8α cells, are TCR-negative intraepithelial lymphocytes (IEL) possessing cytokine and chemokine profiles and functions related to innate immune cells. iCD8α cells constitute an important source of osteopontin in the intestinal epithelium. Osteopontin is a pleiotropic cytokine with diverse roles in bone and tissue remodeling, but also has relevant functions in the homeostasis of immune cells. In this report, we present evidence for the role of iCD8α cells and osteopontin in the homeostasis of TCR-negative NKp46^+^NK1.1^+^ IEL (ILC1-like). We show that in the absence of iCD8α cells, the number of NKp46^+^NK1.1^+^ IEL is significantly reduced. These ILC1-like cells are involved in intestinal pathogenesis in the anti-CD40 mouse model of intestinal inflammation. Reduced iCD8α cell numbers and/or osteopontin expression results in a milder form of intestinal inflammation in this disease model. Collectively, our results suggest that iCD8α cells and osteopontin promote survival of NKp46^+^NK1.1^+^ IEL, which significantly impacts the development of intestinal inflammation.

## Introduction

Intestinal intraepithelial lymphocytes (IEL) constitute a population of cells dwelling interspersed in the monolayer of intestinal epithelial cells (IEC), and represent a unique immunological compartment in the intestines. Because of their anatomical location, IEL are considered to be the first line of defense against the enormous antigenic stimulus present in the lumen of the intestines. T cell receptor αβ^+^ and γδ^+^ cells constitute the great majority of IEL [1–3], and these cells possess many and varied roles during mucosal immune responses and inflammatory processes, ranging from specific immunity against pathogens, tissue repair and homeostasis of the intestinal epithelium [4–9]. Lately, it has been recognized that the IEL compartment also harbors TCR^neg^ lymphoid cells with critical roles in mucosal immune responses [3]. The great majority of TCR^neg^ IEL is composed of cells expressing intracellular CD3γ, which can be divided in CD8αα^+^ or CD8αα^−^ IEL [10]. TCR^neg^CD8αα^+^ IEL, also referred to as innate CD8α (iCD8α) cells, have been previously characterized by our group both in mice and humans [11]. iCD8α cells possess a chemokine and cytokine signature, antigen processing capabilities, and other functions like bacteria uptake, that suggest that these cells are important during early immune responses [11]. Other TCR^neg^ IEL resemble innate lymphoid cells (ILC) with differential expression of the natural cytotoxicity receptor NKp46 [12–14]. Although their function is not completely understood, NKp46^+^NK1.1^+^ IEL have been shown to promote disease development in the anti-CD40 model of colitis [12].

The phosphoprotein osteopontin, encoded by the gene Spp-1, is a glycosylated molecule that was originally characterized as part of the rat bone matrix [15, 16], and later shown to induce Th1 responses, promote pathogenic Th17 survival, enhance NKT cell activation of concanavalin A-induced hepatitis, and regulate the homeostasis and function of NK cells [17-21]. A recent publication shows that lack of osteopontin results in reduced TCRγδ IEL, and that this molecule enhances *in vitro* survival of TCRαβ and TCRγδ IEL [22]. In steady state conditions, iCD8α cells express significant amounts of osteopontin [11], suggesting a potential role for these cells in IEL homeostasis. In terms of intestinal inflammation and disease, osteopontin appears to have divergent roles. For example, in DSS colitis, osteopontin appears to be beneficial during acute disease stages, whereas in chronic disease stages it is detrimental [23]. In trinitrobenzene sulphonic acid-induced colitis, osteopontin enhances development of disease [24]. In humans, plasma osteopontin is increased in individuals with inflammatory bowel diseases (IBD) compared to healthy controls [25, 26]. Although a report indicates that osteopontin is downregulated in the mucosa of Crohn’s disease patients [27], other groups have reported higher osteopontin expression in the intestines of individuals with ulcerative colitis and Crohn’s disease [26, 28]. While these results may be conflicting, they underscore the importance of osteopontin in inflammatory processes and warrant further exploration of this molecule during mucosal immune responses.

In this report we investigated the effect of iCD8α cells as a source of osteopontin in the homeostasis of TCR^neg^ NKp46^+^NK1.1^+^ IEL and their impact in mucosal innate responses. Using mice with reduced iCD8α cell numbers, we show that these cells have a critical role in NKp46^+^NK1.1^+^ IEL survival, which is in part mediated by osteopontin. Disruption of NKp46^+^NK1.1^+^ IEL homeostasis impacts the development of inflammatory processes in the intestines.

## Materials and methods

### Mice

Rag-2^−/−^ mice in the C57BL/6 background have been in our colony for several years; these mice were originally purchased from the Jackson Laboratories. Spp-1^−/−^ mice in the C57BL/6 background were obtained from the Jackson Laboratories. E8_I_^−/−^ mice were graciously provided by Dr. Hilde Cheroutre. Spp-1-GFP-Knock-in mice have been previously reported [22]. To homogenize as much as possible the microbiome, all mice obtained from external sources mice were bred in our facility with Rag-2^−/−^ mice to generate heterozygote mice for both mutations, and from these founders we obtained Spp-1^−/−^Rag-2^−/−^, E8_I_^−/−^Rag-2^−/−^, and Rag-2^−/−^Spp-1-GFP-Knock-in mice. Mice were between 8 to10-week-old. All mice were bred and housed under similar conditions. The Institutional Animal Care and Use Committee at Vanderbilt University Medical Center approved all animal procedures.

### IEL isolation

IEL were isolated by mechanical disruption as previously reported [29]. Briefly, after flushing the intestinal contents with cold HBSS and removing excess mucus, the intestines were cut into small pieces (~1cm long) and shaken for 45 minutes at 37°C in HBSS supplemented with 5% fetal bovine serum and 2mM EDTA. Supernatants were recovered and cells isolated using a discontinuous 40/70% Percoll (General Electric) gradient. In some experiments, IEL preparations were positively enriched using anti-CD45 or anti-CD8α magnetic beads/columns (Miltenyi).

### Reagents and flow cytometry

Fluorochrome-coupled anti-CD8α, -CD45, -NK1.1, and anti-NKp46 were purchased from Thermo Fisher, BD Biosciences or Tombo. Annexin V and 7AAD were purchased from BD Biosciences. All staining samples were acquired using a FACS Canto II Flow System (BD Biosciences) and data analyzed using FlowJo software (Tree Star). Cell staining was performed following conventional techniques. Manufacturer’s instructions were followed for Annexin V staining.

### In vitro survival assay

Enriched CD45^+^ IEL (1×10^5^ cells/well) from Rag-2^−/−^ or E8_I_^−/−^Rag-2^−/−^ mice were cultured in a 96-well flat-bottomed well plate in RPMI complemented with 10% fetal bovine serum, penicillin/streptomycin, HEPES, L-glutamine and β-mercaptoethanol in the presence or absence of 2 μg/ml of recombinant osteopontin (R&D) for 4 hours. After incubation, cells were recovered and stained for surface markers, 7AAD and annexin V. In other experiments, enriched CD45^+^ IEL (1×10^5^ cells/well) from Spp-1^−/−^Rag-2^−/−^ mice were cultured in the presence of enriched iCD8α cells (1×10^5^ cells/well) from Rag-2^−/−^ mice for 4 hours. After incubation, cells were recovered and stained for surface markers, 7AAD and annexin V.

### Induction of intestinal inflammation with anti-CD40 antibodies

Eight to ten-week-old female mice over 18g of weight were treated i.p. with 75 or 150μg of anti-mouse CD40 antibody clone FGK4.5 (Bio X Cell) as previously described [30]. Mice were weighted prior to injection and every day thereafter. Mice were monitored daily for signs of disease such as rectal bleeding, diarrhea and scruffiness. At the end point, a portion of the colon was used for pathological examination and scoring as previously reported [30]. All pathological analysis was performed by a GI pathologist (MBP) in a blind fashion. Some mice were treated with recombinant osteopontin (2 μg per mouse i.p.) or PBS at days −2, −1 and 1 pre- and post-disease induction (75 μg of anti-CD40).

### Real-time PCR

Up to 60 mg of total proximal colon was homogenized using Trizol (Invitrogen) and the RNA was isolated following conventional procedures. RNA was reverse-transcribed using the High Capacity cDNA Transcription Kit (Applied Biosystems). For real-time PCR we used the relative gene expression method [31]. GAPDH served as a normalizer. IL-23p19 primers were purchased from QIAGEN and the sequence for osteopontin primers are: Forward: AGCCACAAGTTTCACAGCCACAAGG; Reverse: CTGAGAAATGAGCAGTTAGTATTCCTGC.

### Osteopontin protein detection

For total osteopontin present in tissue, a ~0.5 cm piece of intestine was cultured in a 24-well plate in RPMI containing 10% fetal bovine serum for 24 hrs at 37°C in 5% CO_2_. Supernatants were collected and cleared. In another experiment, enriched iCD8α cells were cultured at 1×10^5^ cells/well in a 96-well flat-bottomed plate for 24 hr. Osteopontin concentration was determined in the supernatants using a Quantikine ELISA kit (R&D) following manufacturer’s instructions.

### Statistical analysis

Statistical significance between the experimental groups was determined by application of an unpaired two-tailed Student’s t-test or ANOVA using Prism 7. A *p* value <0.05 was considered significant.

## Results

### iCD8α cell deficiency results in decreased NKp46^+^NK1.1^+^ IEL

The study of the innate immune system is facilitated by analyzing mice deficient in adaptive immune cells such as Rag-2^−/−^ mice. Analysis of the IEL compartment in these mice showed two main population of cells present in IEL preparations: a population of large cells composed primarily of IEC, and a population of smaller cells constituting lymphoid cells (Fig. 1a, left dot plot). The latter population consisted primarily of CD45^+^ cells (Fig. 1a, histogram), which could be divided in CD8αα^+^ and CD8αα^neg^ cells (Fig. 1a right dot plots). The former cells constituted iCD8α cells and represented the majority population of innate cells in the IEL compartment of Rag-2^−/−^ mice. Further subdivision of the CD8αα^neg^ cells showed a well-defined population of NKp46^+^NK1.1^+^ IEL, and other IEL with a gradient expression of NK1.1 (Fig. 1a right dot plots). The E8_I_ enhancer region is critical for the expression of CD8α homodimers in lymphoid cells present in the intestinal epithelium, without affecting other cells, such as CD8α^+^ dendritic cells [32, 33]. In a previous publication, we showed that mice deficient in E8_I_ present a significant reduction in iCD8α cells [11], and analysis of E8_I_^−/−^Rag-2^−/−^ mice recapitulated this deficiency (Fig. 1a, right dot plots). Because only iCD8α cells express CD8α homodimers, E8_I_^−/−^ Rag-2^−/−^ mice serve as a model for iCD8α cell deficiency. Interestingly, E8_I_^−/−^Rag-2^−/−^ mice presented with lower numbers of total CD45^+^ IEL, which may account for the reduction in iCD8α cells (Fig. 1b). Moreover, the IEL compartment of E8_I_^−/−^Rag-2^−/−^ mice also presented a significant reduction in the frequencies and cell numbers of NKp46^+^NK1.1^+^ IEL (Fig. 1b). These cells do not express CD8α homodimers (Fig. 1a, right dot plots) and therefore the decrease in numbers is not directly related with the E8_I_ mutation.

**Fig. 1.**
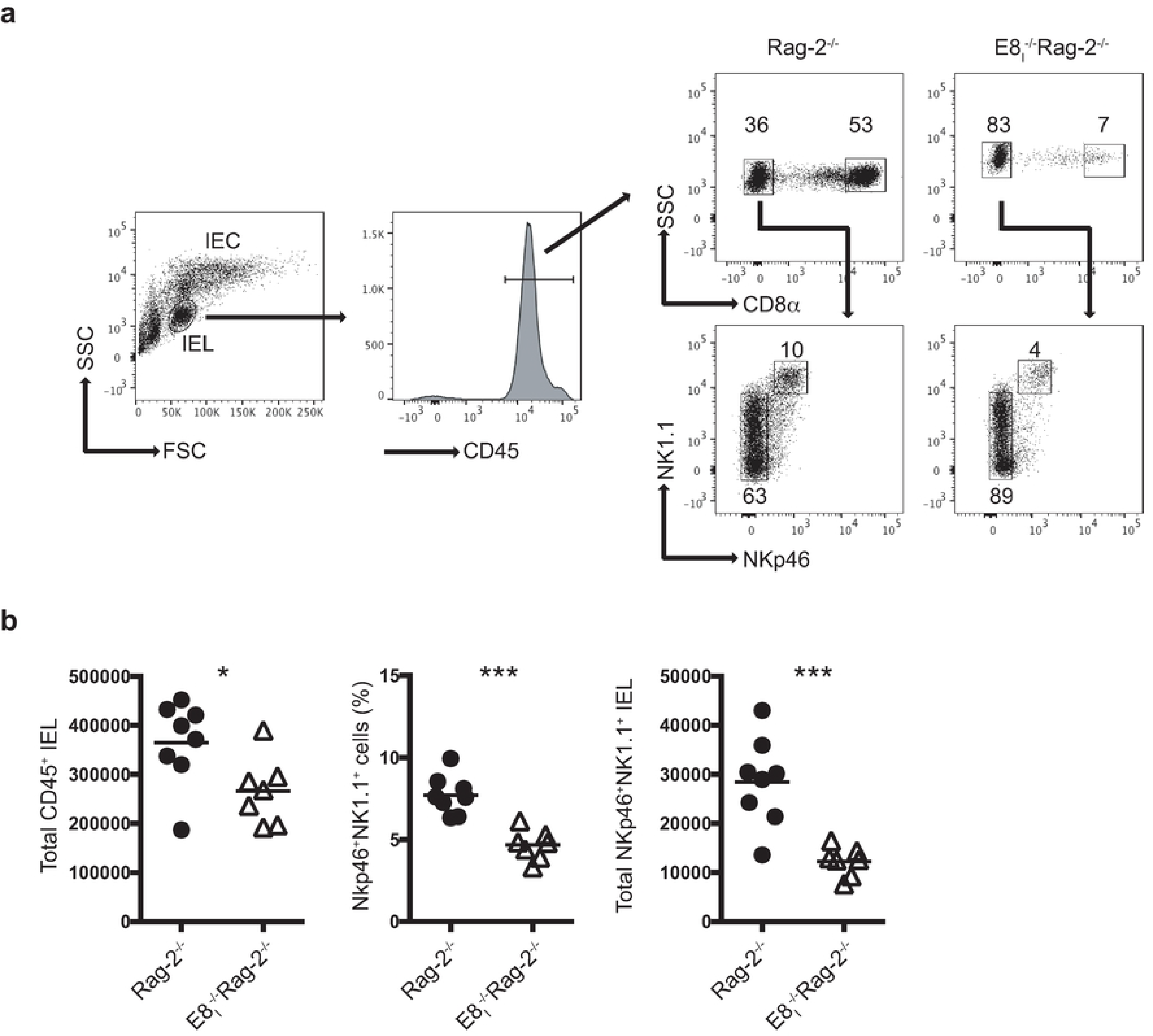
iCD8α cell deficiency results in decreased NKp46^+^NK1.1^+^IEL numbers. Total IEL from Rag-2^−/−^ and E8_I_^−/−^Rag-2^−/−^ mice were analyzed for the presence of CD45^+^ and CD45^+^CD8α^−^ NKp46^+^NK1.1^+^ IEL. (a) Gating strategy for the analysis of the IEL compartment used throughout this report. Dead cells were excluded using a viability dye. (b) Total IEL numbers of the indicated subpopulations. Each symbol represents an individual mouse (n=7 to 8). Data are representative of at least two independent experiments. \**p*<0.05, \*\*\**p*<0.001 using unpaired two-tailed Student’s T test.

### Osteopontin expression in the IEL compartment is primarily associated with iCD8α cells

Osteopontin is a pleiotropic cytokine that has been reported to sustain homeostasis of lymphoid cells, including NK cells [19] and concanavalin A activated T cells [18]. Because iCD8α cells have been reported to be a source of osteopontin, we reasoned that the significant absence of these cells in E8_I_^−/−^Rag-2^−/−^ mice may result in decrease osteopontin production in the intestines. Indeed, the expression of osteopontin mRNA in the intestines of Rag-2^−/−^ mice was significantly higher than that observed in the intestines of E8_I_^−/−^Rag-2^−/−^ mice (Fig. 2a). To investigate osteopontin production in the IEL compartment, we analyzed Rag-2^−/−^ mice carrying the Spp-1-EGFP knock-in reporter gene [22]. Whereas NKp46^+^NK1.1^+^ and other CD8α^−^ IEL (NKp46^−^ NK1.1^lo/−^) presented low GFP staining, most iCD8α cells showed high GFP expression (Fig. 2b), indicating that iCD8α cells are a key source of osteopontin within innate IEL, and corroborate the reduction of this cytokine in mice deficient in iCD8α cells (Fig. 2a).

**Fig. 2.**
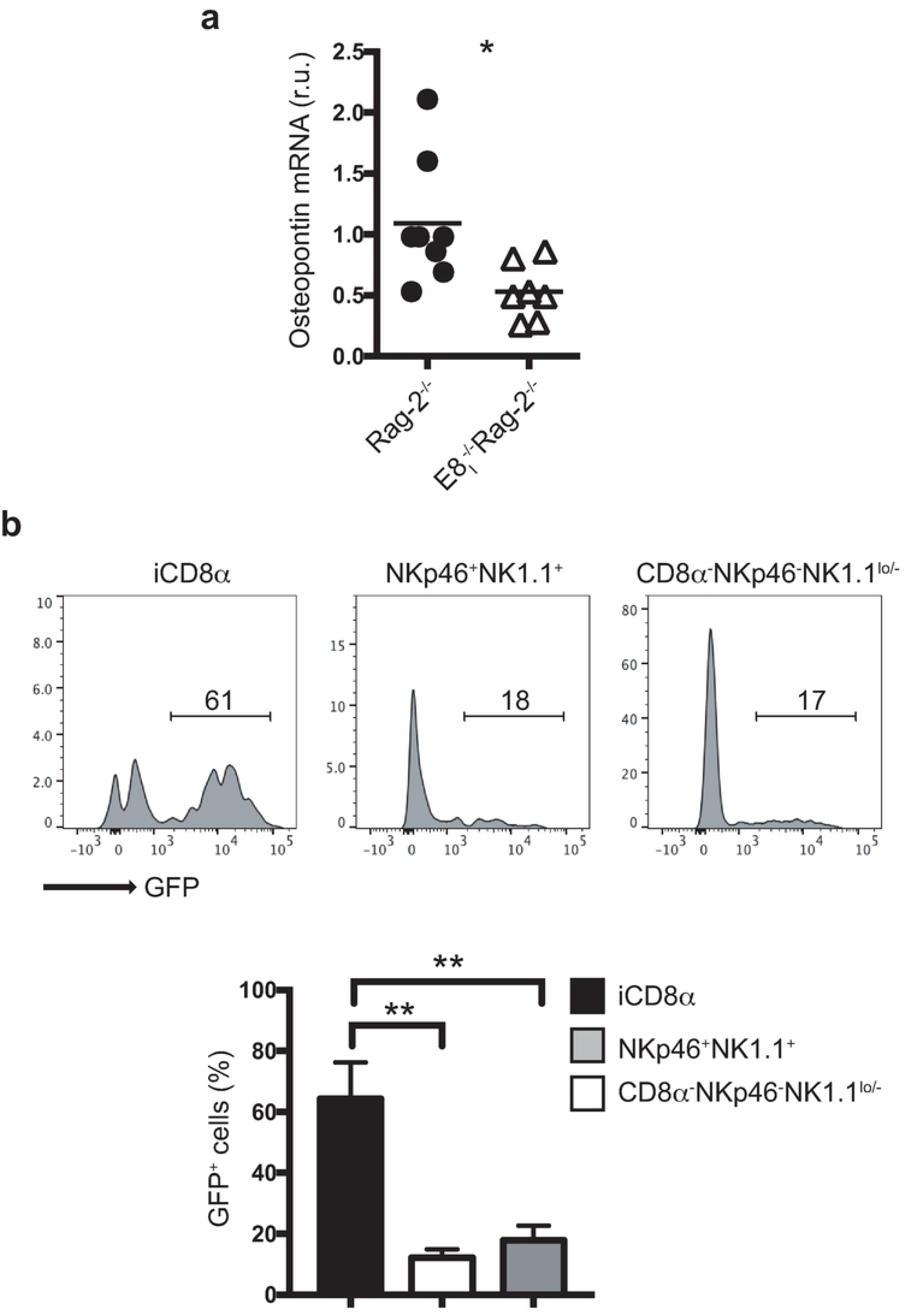
Osteopontin expression in the IEL compartment is primarily associated with iCD8α cells. (a) Osteopontin mRNA expression in colon of the indicated mice. Expression levels in E8_I_^−/−^Rag-2^−/−^ mice were compared to the average expression levels observed in Rag-2^−/−^ mice. Each symbol represents an individual mouse (n=7 to 8). Data are the combination of two independent experiments. (b) Osteopontin expression in naïve Rag-2^−/−^Spp-1-EGFP-KI mice. Histogram is a representative mouse (n=4). Data are representative of at least two independent experiments. \*\**p*<0.01, using unpaired two-tailed Student’s T test for (a) and non-parametric one-way ANOVA for (b).

### iCD8α cells and osteopontin promote survival of NKp46^+^NK1.1^+^ IEL

The above results suggest that iCD8α cell-derived osteopontin is important for maintaining normal levels of NKp46^+^NK1.1^+^ cells. One possibility is that osteopontin promotes the survival of these IEL. To test this hypothesis, total enriched-CD45^+^ IEL from Rag-2^−/−^ mice were cultured for 4 hours in the presence or absence of recombinant osteopontin, and the survival of NKp46^+^NK1.1^+^ IEL was determined by 7AAD and annexin V staining. As shown in Fig. 3a, recombinant osteopontin did not affect annexin V levels in NKp46^+^NK1.1^+^ IEL derived from Rag-2^−/−^ mice, suggesting that osteopontin produced by cells present in the culture (like iCD8α cells, which are the main producers of osteopontin in the intestinal epithelium, Fig.2b) was sufficient to maintain survival, while addition of exogenous osteopontin did not improve survival. However, when enriched CD45^+^ IEL derived from E8_I_^−/−^Rag-2^−/−^ mice (deficient in iCD8α cells) were cultured in the presence of recombinant osteopontin, the levels of annexin V staining were lower than in cells cultured in the absence of recombinant osteopontin (Fig. 3b), which suggests that the addition of osteopontin contributes to the survival of NKp46^+^NK1.1^+^ IEL from E8_I_^−/−^Rag-2^−/−^ mice. To determine the role of iCD8α cells in NKp46^+^NK1.1^+^ IEL survival, CD45^+^ IEL from Spp-1^−/−^Rag-2^−/−^ mice were cultured in the presence or absence of iCD8α cells from Rag-2^−/−^ mice, which produce osteopontin. As seen in Fig. 3c, addition of iCD8α cells decreased the level of annexin V staining in NKp46^+^NK1.1^+^ IEL, indicating that iCD8α cells promote the survival of NKp46^+^NK1.1^+^ IEL.

**Fig. 3.**
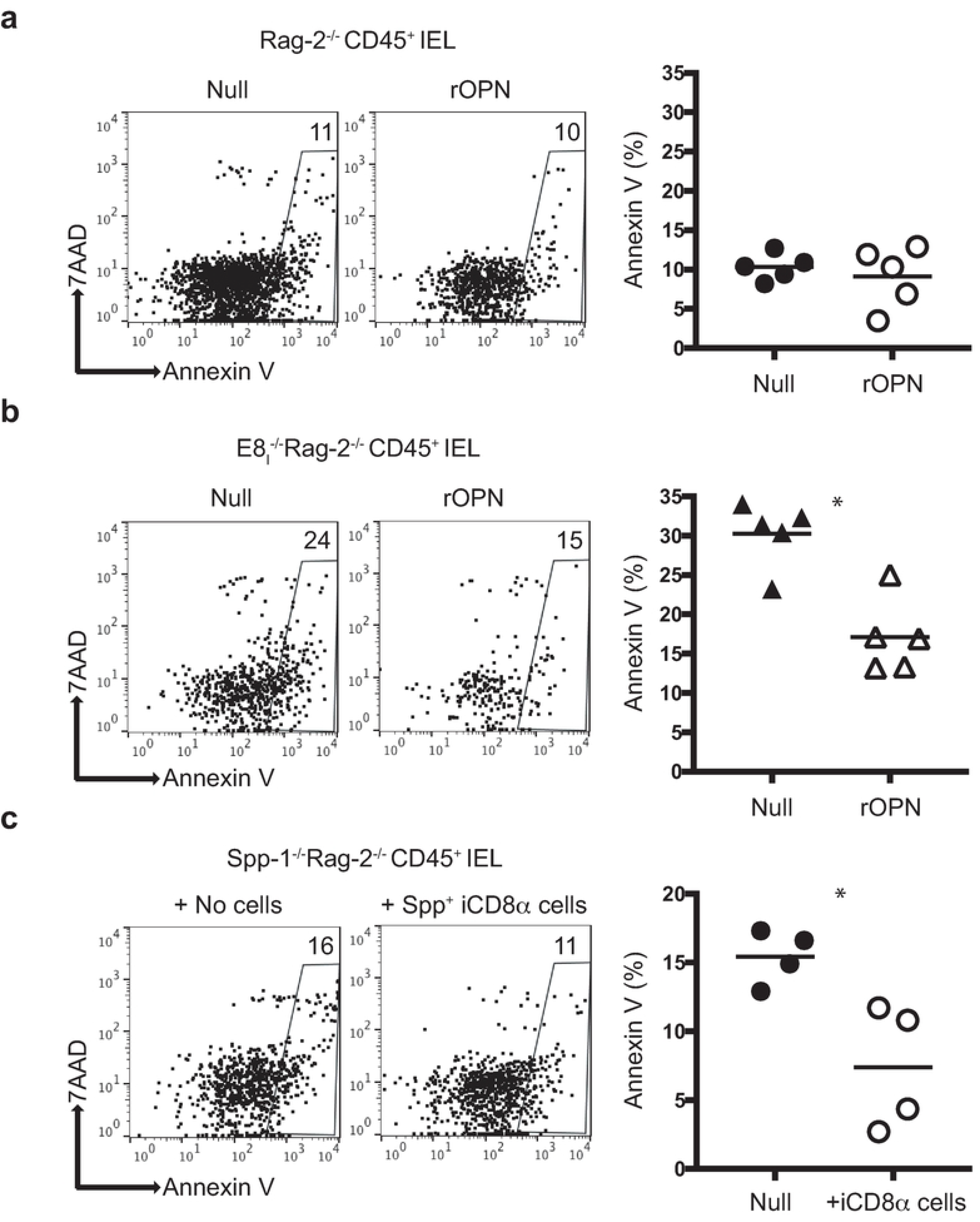
iCD8α cells and osteopontin promote survival of NKp46^+^NK1.1^+^ IEL. Enriched CD45^+^ IEL from Rag-2^−/−^ (a) or E8I^−/−^Rag-2^−/−^ (b) mice were incubated in the presence or absence of recombinant osteopontin (2μg/ml final concentration). Cells were recovered 4 hours later and analyzed for annexin V staining on NKp46^+^NK1.1^+^ IEL as indicated in the Materials and methods section. Data are representative of at least two independent experiments (n=4). (c) Enriched CD45^+^ cells from Spp-1^−/−^Rag-2^−/−^ mice were incubated in the presence or absence of iCD8α cells derived from Rag-2^−/−^ mice. Cells were recovered 4 hours later and analyzed for annexin V staining on NKp46^+^NK1.1^+^ IEL as indicated in the Materials and Method section. In order to obtain enough iCD8α cells, 2-3 mice were pooled and counted as one sample (n=4). Data is representative of at least 2 experiments. \**p*<0.05, using unpaired two-tailed Student’s T test.

Overall, these results indicate that both, iCD8α cells and osteopontin, have an important role in the homeostasis of NKp46^+^NK1.1^+^ IEL.

### Osteopontin kinetics during intestinal inflammation

To investigate the kinetics of osteopontin production during intestinal inflammation, we used the anti-CD40 model of colitis, in which treatment of T and B cell deficient mice (e.g. Rag-2^−/−^) with anti-CD40 results in weight loss, loose stools, rectal bleeding and inflammation of the colon mediated by IL-23 [30]. This system represents a good model for the analysis of innate immune responses during intestinal inflammation. We treated Rag-2^−/−^ mice with anti-CD40 and 2 days after, osteopontin protein levels were measured in colon tissue or enriched-iCD8α cells. Osteopontin was readily detected either in total colon (Fig. 4a) or iCD8α cells (Fig. 4b) derived from anti-CD40-treated mice in comparison to naïve animals. To determine the kinetics of osteopontin expression in the intestinal epithelium during inflammation, we treated Rag-2^−/−^Spp-1-EGFP knock-in reporter mice with anti-CD40. The expression of osteopontin in iCD8α cells remained constant 3-and 7-days after disease induction, whereas expression of osteopontin in NKp46^+^NK1.1^+^ IEL increased at 3- and 7-days post-treatment (Fig. 4c). On the other hand, expression of osteopontin in other IEL populations (represented as CD8α^−^NKp46^−^NK1.1^lo/−^) decreased during the course of the disease (Fig. 4c). These results indicate that during anti-CD40-induced colitis, iCD8α cells, and to a lesser extent NKp46^+^NK1.1^+^ IEL comprise significant sources of osteopontin in the intestinal epithelium.

**Fig. 4.**
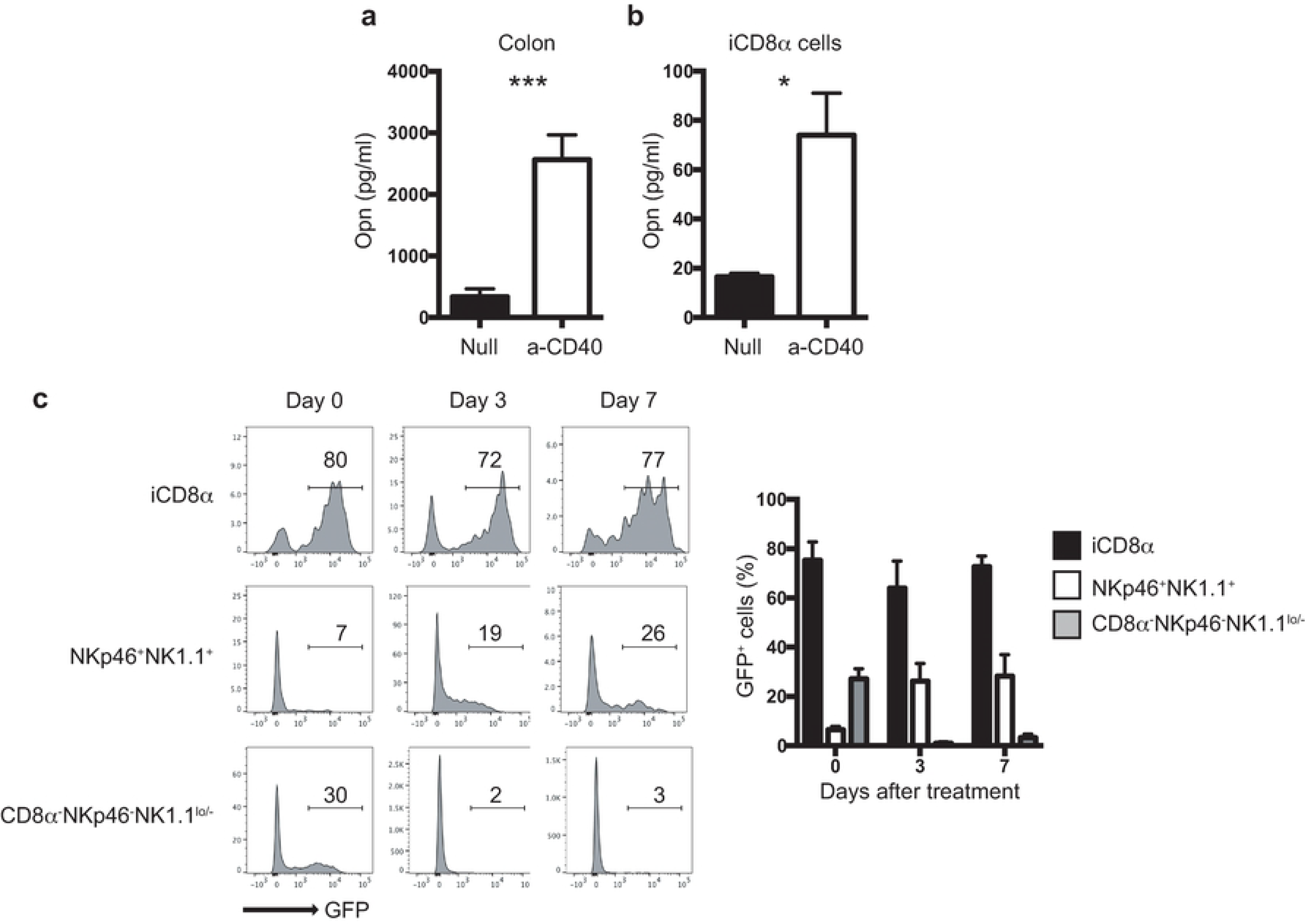
Osteopontin kinetics during intestinal inflammation. (a) Osteopontin protein concentration in the supernatants of whole colon tissue cultures from naïve and anti-CD40-treated Rag-2^−/−^ mice. Data is representative of at least 2 experiments (n=5). (b) Osteopontin protein concentration in the supernatants of iCD8α cells derived from naïve and anti-CD40-treated Rag-2^−/−^ mice. Data is representative of at least two experiments. In order to obtain enough iCD8α cells, 2-3 mice were pooled and counted as one sample (n=3). (c) GFP expression in Rag-2^−/−^Spp-1-EGFP-KI mice treated with anti-CD40 and analyzed at the indicated time points. Cells were gated as indicated in Fig.1a. Histograms are from a representative sample. Bar graph shows data summary. \**p*<0.05, \*\*\**p*<0.001 using unpaired two-tailed Student’s T test.

### Osteopontin promotes intestinal inflammation

The increase in osteopontin production observed during anti-CD40-induced colitis suggests and important role for this cytokine in disease development. To test this hypothesis, we treated Rag-2^−/−^ and osteopontin-deficient Rag-2^−/−^ (Spp-1^−/−^Rag-2^−/−^) mice with anti-CD40 and monitored the mice for 7 days. As expected, Rag-2^−/−^ mice lost weight starting at day 1 post treatment, which was also observed in Spp-1^−/−^Rag-2^−/−^ mice (Fig. 5a). However, after day 2, Spp-1^−/−^Rag-2^−/−^ mice showed significantly decreased weight loss in comparison to Rag-2^−/−^ mice, and presented less colon pathology at the end of the experiment (Fig. 5b). The decrease in disease observed was accompanied by reduced levels of IL-23 expression in the colon (Fig. 5c). These results indicate that osteopontin is detrimental in this model of intestinal inflammation.

**Fig. 5.**
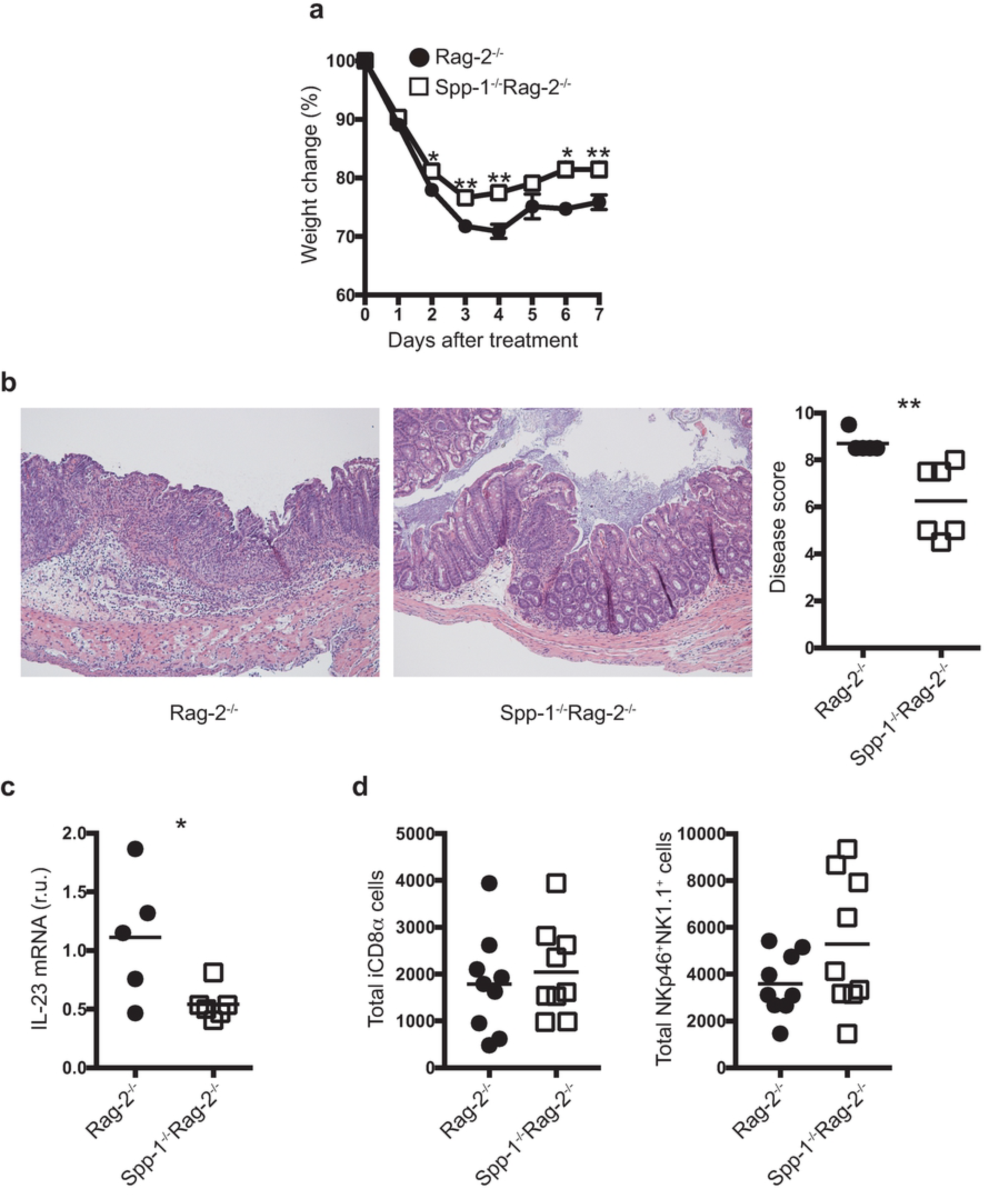
Osteopontin promotes intestinal inflammation. (a) Rag-2^−/−^ and Spp-1^−/−^Rag-2^−/−^ mice were treated with 150 μg of anti-CD40 and monitored daily for weight change for 7 days. (b) Pathological representation (micrographs, magnification 200X) and disease score; each symbol represents an individual mouse (n=5 to 6). Data is representative of at least 2 experiments. (c) IL-23p19 mRNA expression from total colons; each symbol represents an individual mouse (n=5 to 6). Data is representative of at least 2 experiments. (d) Total iCD8α cell and NKp46^+^NK1.1^+^ IEL numbers in naïve Rag-2^−/−^ and Spp-1^−/−^Rag-2^−/−^ mice; each symbol represents and individual mouse (n=9 to 10). Data is representative of 3 experiments. \**p*<0.05, \*\**p*<0.01 using two-way ANOVA (a) or unpaired two-tailed Student’s T test (b, c, d).

Interestingly, the numbers of iCD8α cells and NKp46^+^NK1.1^+^ IEL in naïve Spp-1^−/−^Rag-2^−/−^ mice were comparable to those observed in Rag-2^−/−^ animals (Fig. 5d), suggesting that the absence of osteopontin has a greater influence in disease development than in the reduction of NKp46^+^NK1.1^+^ IEL numbers.

### Decreased intestinal inflammation in mice deficient in iCD8a cells

To investigate whether iCD8α IEL deficiency has an impact in intestinal inflammation, we treated E8_I_^−/−^Rag-2^−/−^ mice and control Rag-2^−/−^ mice with anti-CD40. E8_I_^−/−^Rag-2^−/−^ mice lost less weight throughout the course of the experiment (Fig. 6a) and presented less colon pathology (Fig. 6b), mirroring the observed results in Spp-1^−/−^Rag-2^−/−^ mice (Fig. 5a). Analysis of the kinetics of osteopontin expression in the colon of anti-CD40-treated Rag-2^−/−^ and E8_I_^−/−^Rag-2^−/−^ mice showed incremental expression of osteopontin mRNA at day 2 and 7 post disease induction; however, the levels of osteopontin mRNA levels were consistently lower in E8_I_^−/−^Rag-2^−/−^ mice than in Rag-2^−/−^ mice (Fig. 6c). To investigate whether treatment with osteopontin increases disease severity in iCD8α cell-deficient mice, E8_I_^−/−^Rag-2^−/−^ mice were injected i.p. at day −2, −1, and 0 with recombinant osteopontin, followed by disease induction with a reduced dose of anti-CD40 at day 0 (a lower dose was chosen to better detect changes in disease severity). Rag-2^−/−^ control mice lost similar weight with low anti-CD40 than mice treated with the regular dose (compare Fig. 6a and 6d); however, E8_I_^−/−^Rag-2^−/−^ mice treated with low anti-CD40 recovered faster than E8_I_^−/−^Rag-2^−/−^ mice treated with the full anti-CD40 dose (compare Fig. 6a and 6d). Although, E8_I_^−/−^Rag-2^−/−^ mice treated with recombinant osteopontin presented weight loss similar to PBS-treated E8_I_^−/−^Rag-2^−/−^ mice during the first few days after disease induction, the former group did not recover as the PBS-treated group and their weights were more similar to Rag-2^−/−^ control mice. (Fig. 6d). Although colon pathological scores were comparable between the control and recombinant osteopontin-treated E8_I_^−/−^Rag-2^−/−^ groups, there was a tendency for higher disease severity in the latter group. Therefore, our results indicate that administration of osteopontin slightly increases disease severity in the absence of iCD8α cells and low osteopontin expression in the colon.

**Fig. 6.**
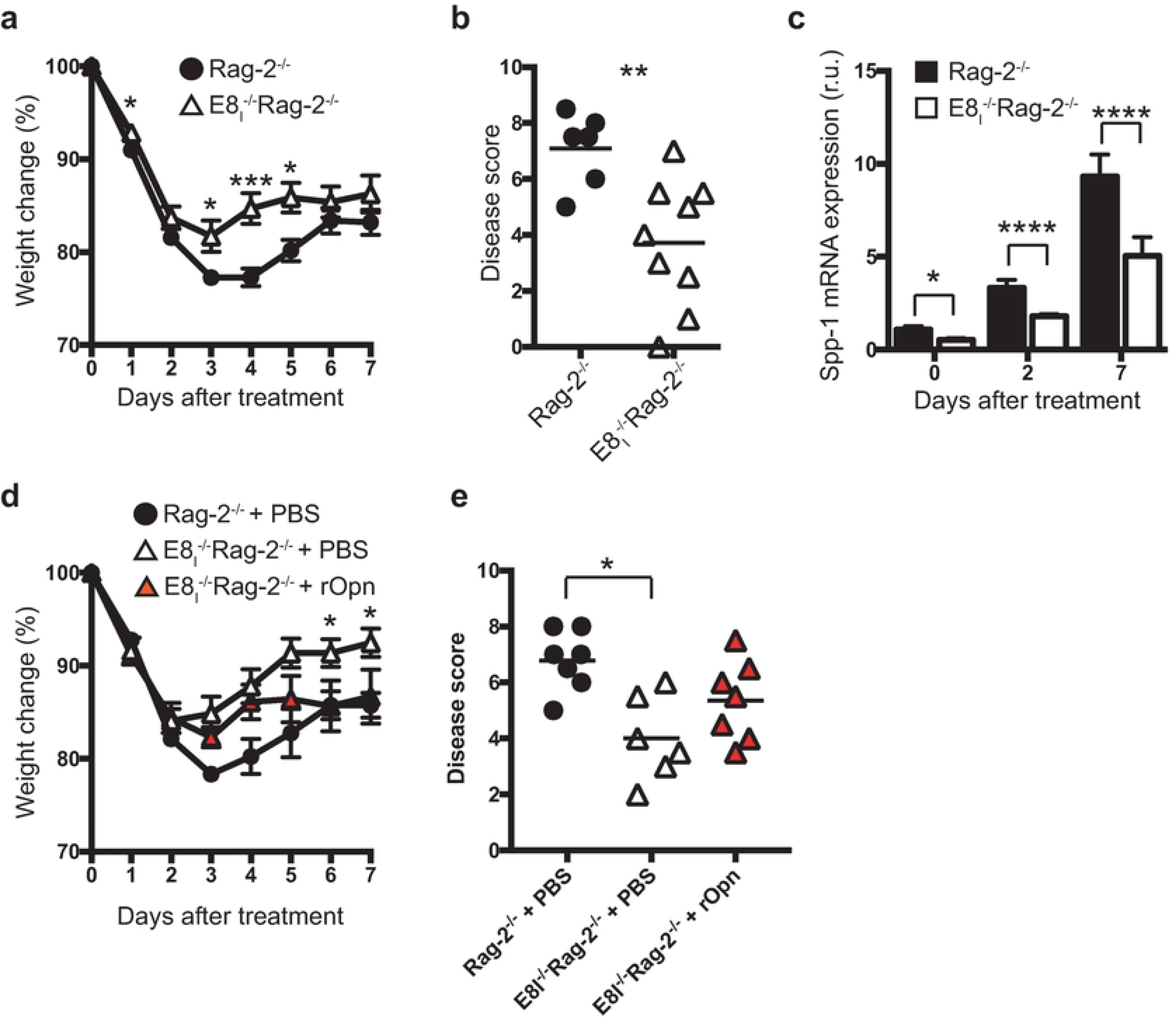
Decreased intestinal inflammation in mice deficient in iCD8α cells. Rag-2^−/−^ and E8_I_^−/−^ Rag-2^−/−^ mice were treated with anti-CD40 and monitored for 7 days for weight change (a). (b) At the endpoint, colons were harvested for pathological analysis. (c) Osteopontin mRNA expression from the colons of anti-CD40 treated Rag-2^−/−^ and E8_I_^−/−^Rag-2^−/−^ mice at the indicated time points. Data is representative of at least two independent experiments (n=6 to 8). (d) E8_I_^−/−^Rag-2^−/−^ mice were treated with recombinant osteopontin or PBS at day −2, −1 and 1 before and after disease induction with 70μg of anti-CD40 antibodies, and their weights monitored for 7 days. (e) At the endpoint, colons were harvested for pathological analysis. Data is representative of at least 3 independent experiments (n=6 to7). Each symbol represents and individual mouse. \**p*<0.05, \*\**p*<0.01, ***p<0.01 using two-way ANOVA (a, d), unpaired two-tailed Student’s T test (b, c) or one-way ANOVA (e).

## Discussion

Osteopontin is known to be widely expressed in the intestinal mucosa of ulcerative colitis and Crohn’s disease patients, and in the latter group, osteopontin plasma levels are increased in comparison to control individuals [26, 34], suggesting an involvement of this molecule in the pathology of inflammatory bowel diseases. However, the role of osteopontin in mouse models of intestinal inflammation is controversial. In the DSS model of colitis, reports vary about the role of osteopontin, either as a pro or anti-inflammatory factor [23, 34-36]. Moreover, in the trinitrobenzene sulphonic acid-induced model of colitis, osteopontin-deficient mice fare better than wild type animals, suggesting a pro-inflammatory role for this cytokine [37]. In contrast, in the IL-10-deficiency model of spontaneous intestinal inflammation, IL-10^−/−^Spp-1^−/−^ mice develop disease faster than IL-10^−/−^ control mice [36]. Finally, adoptive transfer of naïve CD62L^hi^CD4^+^ T cells into Rag-2^−/−^Spp-1^−/−^ mice resulted in less chronic colitis than Rag-2^−/−^ recipient mice [38]. How to reconcile these diverse observations? It is possible that osteopontin’s impact varies depending on the primary cell populations responsible for disease induction or the disease stage. For example, here we propose that iCD8α cells, via osteopontin, promote the survival of pro-inflammatory NKp46^+^NK1.1^+^ IEL during acute colitis, whereas in other disease models osteopontin may differentially impact acute and chronic inflammation [23]. Therefore, dissecting how osteopontin affects different branches of the mucosal immune system during steady state levels and inflammatory processes is of critical relevance to increase our understanding of IEL biology and osteopontin function, as well as the impact of this cytokine in diseases such as ulcerative colitis and Crohn’s disease.

IEL reside in a unique anatomical location intercalated between IEC, and in close proximity to the contents of the intestinal lumen. This location makes a unique niche for IEL. In this environment, IEL are most likely subjected to distinctive signals during steady-state levels as well as during intestinal immune responses. In addition, IEL represent a heterogeneous population of lymphocytes with different developmental origins and immunological roles [1–3], and because of this diversity, each IEL population may be subjected to particular environmental clues. How different IEL populations survive and maintain homeostasis in the intestinal epithelium is not very well understood. In this report, we examine the role of a novel IEL population referred to as iCD8α cells and the pleiotropic cytokine osteopontin in the homeostasis of NKp46^+^NK1.1^+^ IEL. iCD8α cells promote clearance of the colitis-inducing pathogen *Citrobacter rodentium* [11], but also exacerbate colitis via granzymes when not properly regulated [39]. The results presented in this report add a new role for iCD8α IEL as a population promoting the survival of NKp46^+^NK1.1^+^ IEL via osteopontin.

Although most of the IEL studies have primarily focused on TCR^+^ IEL (TCRαβ and γδ), recently it has become apparent that TCR^neg^ IEL constitute an important fraction of the IEL compartment. Three distinct TCR^neg^ IEL populations have been characterized to date: iCD3^+^, iCD8α, and ILC-like IEL [10, 12-14]. iCD3^+^ and iCD8α cells appear to be related IEL populations that require IL-15 for their development. How the homeostasis of these cells is maintained in the intestinal epithelium is not clearly understood. There is evidence suggesting that the thymus leukemia (TL) antigen, a ligand for CD8αα homodimers [40, 41], is needed for maintenance of iCD8α cells [11]. Some ILC-like IEL require IL-15 for their survival, such as NKp46-negative IEL [14]. Although osteopontin deficient mice present similar numbers of NKp46^+^NK1.1^+^ IEL (Fig. 5d), their numbers are significantly decreased in mice deficient in iCD8α cells and reduced intestinal osteopontin (E8_I_^−/−^Rag-2^−/−^ mice). These observations raise the possibility that iCD8α cells support NKp46^+^NK1.1^+^ IEL homeostasis by unknown mechanisms in addition to osteopontin.

Our *in vivo* evidence presented in this report indicates that iCD8α cells represent one of the innate IEL populations with highest levels of osteopontin expression, and that mice deficient in iCD8α cells also present decreased osteopontin levels in the colon (Fig. 2a). These results suggest a putative role for iCD8α cells as a source of osteopontin in the intestinal epithelium, allowing proper survival of other IEL in steady state conditions. Although NKp46^+^NK1.1^+^ IEL do not produce osteopontin during steady state conditions, the expression of this cytokine incrementally appears at day 3 and 7 post anti-CD40 treatment. At this moment, the significance of NKp46^+^NK1.1^+^ IEL-derived osteopontin during inflammation is unknown.

It is important to mention that in order to study innate IEL, our results are based on mice lacking TCR^+^ IEL, and therefore, we do not discard the possibility that some TCR^+^ IEL may be osteopontin producers in wild type mice. Indeed, in steady state conditions, using an osteopontin-GFP reporter system, Hattori’s group showed that TCR^+^CD8α^+^ IEL represent a source of osteopontin in the intestines of wild type mice [22]. This group also showed that TCRγδ^+^ IEL *in vivo* are dependent on osteopontin for their survival, whereas in *in vitro* conditions, both TCRαβ^+^ and γδ^+^ IEL survival is blunted by anti-osteopontin antibodies. Although, the report by Hattori’s group and our results presented herein clearly indicate an important role for osteopontin in IEL survival, there is still a significant gap in knowledge about the homeostasis of IEL subpopulations, such as TCRβ^+^CD4^+^, TCRβ^+^CD4^+^CD8αα^+^, TCRβ^+^CD8αβ^+^, TCRβ^+^CD8αα^+^ and CD8αα^neg^iCD3^+^ cells; similarly, it is unknown whether human IEL require osteopontin for their survival/homeostasis. Another outstanding question is the receptor used by osteopontin to stimulate IEL. One possible candidate is CD44, a molecule expressed in activated T cells, with the capacity of binding osteopontin [42].

It is poorly investigated whether different population of IEL interact with each other. There are few reports that indirectly suggest that this could be the case, for example, TCRγδ IEL control the activation status and numbers of TCRαβ^+^CD8αβ^+^ IEL in humans [43], whereas iCD8α cells may present antigen to CD4^+^ IEL in an MHC class II restricted fashion [11]. Although our results do not provide direct evidence showing interaction between iCD8α cells and NKp46^+^NK1.1^+^ IEL in the intestinal epithelium, IEL may either directly interact with each other or may communicate via cytokines and/or other factors. However, more research needs to be done to have a better understanding of IEL-IEL interactions.

In conclusion, in this report we provide evidence indicating an important and novel role for iCD8α cells in the homeostasis of NKp46^+^NK1.1^+^ IEL. We also show that the effect of iCD8α cells is mediated in part by osteopontin, which adds to the growing roles of this cytokine in different biological processes.

## Acknowledgements

We thank the Translational Pathology Shared Resource for tissue processing. This work was supported by NIH grant R01DK111671 (to D.O-V.); Careers in Immunology Fellowship Program from the American Association of Immunologist (to D.O-V. and A.N.); National Library of Medicine T15 LM00745 grant (to M.J.G.); and scholarships from the Digestive Disease Research Center at Vanderbilt University Medical Center supported by NIH grant P30DK058404.

## References

1. Van Kaer L, Olivares-Villagomez D. Development, Homeostasis, and Functions of Intestinal Intraepithelial Lymphocytes. J Immunol. 2018;200(7):2235–44. Epub 2018/03/21. doi: 10.4049/jimmunol.1701704. PubMed PMID: 29555677; PubMed Central PMCID: PMCPMC5863587.

2. Cheroutre H. Starting at the beginning: new perspectives on the biology of mucosal T cells. Annu Rev Immunol. 2004;22:217–46. PubMed PMID: 15032579.

3. Olivares-Villagomez D, Van Kaer L. Intestinal Intraepithelial Lymphocytes: Sentinels of the Mucosal Barrier. Trends Immunol. 2017. Epub 2017/12/10. doi: 10.1016/j.it.2017.11.003. PubMed PMID: 29221933.

4. Chen Y, Chou K, Fuchs E, Havran WL, Boismenu R. Protection of the intestinal mucosa by intraepithelial gamma delta T cells. Proc Natl Acad Sci U S A. 2002;99(22):14338–43. PubMed PMID: 12376619.

5. Das G, Augustine MM, Das J, Bottomly K, Ray P, Ray A. An important regulatory role for CD4+CD8 alpha alpha T cells in the intestinal epithelial layer in the prevention of inflammatory bowel disease. Proc Natl Acad Sci U S A. 2003;100(9):5324–9. PMC1535703. PubMed PMID: 12695566.

6. Guy-Grand D, DiSanto JP, Henchoz P, Malassis-Seris M, Vassalli P. Small bowel enteropathy: role of intraepithelial lymphocytes and of cytokines (IL-12, IFN-gamma, TNF) in the induction of epithelial cell death and renewal. Eur J Immunol. 1998;28(2):730–44. PubMed PMID: 9521083.

7. Inagaki-Ohara K, Chinen T, Matsuzaki G, Sasaki A, Sakamoto Y, Hiromatsu K, et al. Mucosal T cells bearing TCRgammadelta play a protective role in intestinal inflammation. J Immunol. 2004;173(2):1390–8. PubMed PMID: 15240735.

8. Lepage AC, Buzoni-Gatel D, Bout DT, Kasper LH. Gut-derived intraepithelial lymphocytes induce long term immunity against Toxoplasma gondii. J Immunol. 1998;161(9):4902–8. PubMed PMID: 9794424.

9. Masopust D, Jiang J, Shen H, Lefrancois L. Direct analysis of the dynamics of the intestinal mucosa CD8 T cell response to systemic virus infection. J Immunol. 2001;166(4):2348–56. PubMed PMID: 11160292.

10. Ettersperger J, Montcuquet N, Malamut G, Guegan N, Lopez-Lastra S, Gayraud S, et al. Interleukin-15-Dependent T-Cell-like Innate Intraepithelial Lymphocytes Develop in the Intestine and Transform into Lymphomas in Celiac Disease. Immunity. 2016;45(3):610–25. Epub 2016/09/11. doi: 10.1016/j.immuni.2016.07.018. PubMed PMID: 27612641.

11. Van Kaer L, Algood HM, Singh K, Parekh VV, Greer MJ, Piazuelo MB, et al. CD8alphaalpha(+) Innate-Type Lymphocytes in the Intestinal Epithelium Mediate Mucosal Immunity. Immunity. 2014;41(3):451–64. Epub 2014/09/16. doi: 10.1016/j.immuni.2014.08.010. PubMed PMID: 25220211; PubMed Central PMCID: PMC4169715.

12. Fuchs A, Vermi W, Lee JS, Lonardi S, Gilfillan S, Newberry RD, et al. Intraepithelial type 1 innate lymphoid cells are a unique subset of IL-12- and IL-15-responsive IFN-gamma-producing cells. Immunity. 2013;38(4):769–81. Epub 2013/03/05. doi: 10.1016/j.immuni.2013.02.010. PubMed PMID: 23453631; PubMed Central PMCID: PMC3634355.

13. Talayero P, Mancebo E, Calvo-Pulido J, Rodriguez-Munoz S, Bernardo I, Laguna-Goya R, et al. Innate Lymphoid Cells Groups 1 and 3 in the Epithelial Compartment of Functional Human Intestinal Allografts. Am J Transplant. 2016;16(1):72–82. Epub 2015/09/01. doi: 10.1111/ajt.13435. PubMed PMID: 26317573.

14. Van Acker A, Gronke K, Biswas A, Martens L, Saeys Y, Filtjens J, et al. A Murine Intestinal Intraepithelial NKp46-Negative Innate Lymphoid Cell Population Characterized by Group 1 Properties. Cell Rep. 2017;19(7):1431–43. doi: 10.1016/j.celrep.2017.04.068. PubMed PMID: 28514662.

15. Franzen A, Heinegard D. Isolation and characterization of two sialoproteins present only in bone calcified matrix. Biochem J. 1985;232(3):715–24. Epub 1985/12/15. PubMed PMID: 4091817; PubMed Central PMCID: PMCPMC1152943.

16. Prince CW, Oosawa T, Butler WT, Tomana M, Bhown AS, Bhown M, et al. Isolation, characterization, and biosynthesis of a phosphorylated glycoprotein from rat bone. J Biol Chem. 1987;262(6):2900–7. Epub 1987/02/25. PubMed PMID: 3469201.

17. Ashkar S, Weber GF, Panoutsakopoulou V, Sanchirico ME, Jansson M, Zawaideh S, et al. Eta-1 (osteopontin): an early component of type-1 (cell-mediated) immunity. Science. 2000;287(5454):860–4. Epub 2000/02/05. PubMed PMID: 10657301.

18. Hur EM, Youssef S, Haws ME, Zhang SY, Sobel RA, Steinman L. Osteopontin-induced relapse and progression of autoimmune brain disease through enhanced survival of activated T cells. Nat Immunol. 2007;8(1):74–83. Epub 2006/12/05. doi: 10.1038/ni1415. PubMed PMID: 17143274.

19. Leavenworth JW, Verbinnen B, Wang Q, Shen E, Cantor H. Intracellular osteopontin regulates homeostasis and function of natural killer cells. Proc Natl Acad Sci U S A. 2015;112(2):494–9. Epub 2015/01/01. doi: 10.1073/pnas.1423011112. PubMed PMID: 25550515; PubMed Central PMCID: PMC4299239.

20. Murugaiyan G, Mittal A, Weiner HL. Increased osteopontin expression in dendritic cells amplifies IL-17 production by CD4+ T cells in experimental autoimmune encephalomyelitis and in multiple sclerosis. J Immunol. 2008;181(11):7480–8. Epub 2008/11/20. PubMed PMID: 19017937; PubMed Central PMCID: PMC2653058.

21. Staines KA, MacRae VE, Farquharson C. The importance of the SIBLING family of proteins on skeletal mineralisation and bone remodelling. The Journal of endocrinology. 2012;214(3):241–55. Epub 2012/06/16. doi: 10.1530/JOE-12-0143. PubMed PMID: 22700194.

22. Ito K, Nakajima A, Fukushima Y, Suzuki K, Sakamoto K, Hamazaki Y, et al. The potential role of Osteopontin in the maintenance of commensal bacteria homeostasis in the intestine. PLoS One. 2017;12(3):e0173629. Epub 2017/03/16. doi: 10.1371/journal.pone.0173629. PubMed PMID: 28296922; PubMed Central PMCID: PMCPMC5351998.

23. Heilmann K, Hoffmann U, Witte E, Loddenkemper C, Sina C, Schreiber S, et al. Osteopontin as two-sided mediator of intestinal inflammation. Journal of cellular and molecular medicine. 2009;13(6):1162–74. Epub 2008/07/17. doi: 10.1111/j.1582-4934.2008.00428.x. PubMed PMID: 18627421; PubMed Central PMCID: PMC4496111.

24. Oz HS, Zhong J, de Villiers WJ. Osteopontin ablation attenuates progression of colitis in TNBS model. Dig Dis Sci. 2012;57(6):1554–61. Epub 2011/12/17. doi: 10.1007/s10620-011-2009-z. PubMed PMID: 22173746.

25. Mishima R, Takeshima F, Sawai T, Ohba K, Ohnita K, Isomoto H, et al. High plasma osteopontin levels in patients with inflammatory bowel disease. J Clin Gastroenterol. 2007;41(2):167–72. Epub 2007/01/25. doi: 10.1097/MCG.0b013e31802d6268. PubMed PMID: 17245215.

26. Sato T, Nakai T, Tamura N, Okamoto S, Matsuoka K, Sakuraba A, et al. Osteopontin/Eta-1 upregulated in Crohn’s disease regulates the Th1 immune response. Gut. 2005;54(9):1254–62. Epub 2005/08/16. doi: 10.1136/gut.2004.048298. PubMed PMID: 16099792; PubMed Central PMCID: PMC1774642.

27. Gassler N, Autschbach F, Gauer S, Bohn J, Sido B, Otto HF, et al. Expression of osteopontin (Eta-1) in Crohn disease of the terminal ileum. Scand J Gastroenterol. 2002;37(11):1286–95. Epub 2002/12/06. PubMed PMID: 12465727.

28. Masuda H, Takahashi Y, Asai S, Takayama T. Distinct gene expression of osteopontin in patients with ulcerative colitis. J Surg Res. 2003;111(1):85–90. Epub 2003/07/05. PubMed PMID: 12842452.

29. Olivares-Villagomez D, Mendez-Fernandez YV, Parekh VV, Lalani S, Vincent TL, Cheroutre H, et al. Thymus leukemia antigen controls intraepithelial lymphocyte function and inflammatory bowel disease. Proc Natl Acad Sci U S A. 2008;105(46):17931–6. PMC2584730. PubMed PMID: 19004778.

30. Uhlig HH, McKenzie BS, Hue S, Thompson C, Joyce-Shaikh B, Stepankova R, et al. Differential activity of IL-12 and IL-23 in mucosal and systemic innate immune pathology. Immunity. 2006;25(2):309–18. Epub 2006/08/22. doi: 10.1016/j.immuni.2006.05.017. PubMed PMID: 16919486.

31. Giulietti A, Overbergh L, Valckx D, Decallonne B, Bouillon R, Mathieu C. An overview of real-time quantitative PCR: applications to quantify cytokine gene expression. Methods. 2001;25(4):386–401. Epub 2002/02/16. doi: 10.1006/meth.2001.1261. PubMed PMID: 11846608.

32. Ellmeier W, Sawada S, Littman DR. The regulation of CD4 and CD8 coreceptor gene expression during T cell development. Annu Rev Immunol. 1999;17:523–54. Epub 1999/06/08. doi: 10.1146/annurev.immunol.17.1.523. PubMed PMID: 10358767.

33. Taniuchi I, Ellmeier W, Littman DR. The CD4/CD8 lineage choice: new insights into epigenetic regulation during T cell development. Advances in immunology. 2004;83:55–89. Epub 2004/05/12. doi: 10.1016/S0065-2776(04)83002-5. PubMed PMID: 15135628.

34. Masuda H, Takahashi Y, Asai S, Hemmi A, Takayama T. Osteopontin expression in ulcerative colitis is distinctly different from that in Crohn’s disease and diverticulitis. J Gastroenterol. 2005;40(4):409–13. Epub 2005/05/04. doi: 10.1007/s00535-005-1567-2. PubMed PMID: 15868372.

35. Zhong J, Eckhardt ER, Oz HS, Bruemmer D, de Villiers WJ. Osteopontin deficiency protects mice from Dextran sodium sulfate-induced colitis. Inflamm Bowel Dis. 2006;12(8):790–6. Epub 2006/08/19. PubMed PMID: 16917234.

36. Toyonaga T, Nakase H, Ueno S, Matsuura M, Yoshino T, Honzawa Y, et al. Osteopontin Deficiency Accelerates Spontaneous Colitis in Mice with Disrupted Gut Microbiota and Macrophage Phagocytic Activity. PLoS One. 2015;10(8):e0135552. Epub 2015/08/15. doi: 10.1371/journal.pone.0135552. PubMed PMID: 26274807; PubMed Central PMCID: PMC4537118.

37. Kourepini E, Aggelakopoulou M, Alissafi T, Paschalidis N, Simoes DC, Panoutsakopoulou V. Osteopontin expression by CD103-dendritic cells drives intestinal inflammation. Proc Natl Acad Sci U S A. 2014;111(9):E856–65. Epub 2014/02/20. doi: 10.1073/pnas.1316447111. PubMed PMID: 24550510; PubMed Central PMCID: PMC3948306.

38. Kanayama M, Xu S, Danzaki K, Gibson JR, Inoue M, Gregory SG, et al. Skewing of the population balance of lymphoid and myeloid cells by secreted and intracellular osteopontin. Nat Immunol. 2017;18(9):973–84. Epub 2017/07/04. doi: 10.1038/ni.3791. PubMed PMID: 28671690; PubMed Central PMCID: PMCPMC5568448.

39. Kumar AA, Delgado AG, Piazuelo MB, Van Kaer L, Olivares-Villagomez D. Innate CD8alphaalpha+ lymphocytes enhance anti-CD40 antibody-mediated colitis in mice. Immun Inflamm Dis. 2017;5(2):109–23. doi: 10.1002/iid3.146. PubMed PMID: 28474503; PubMed Central PMCID: PMCPMC5418141.

40. Leishman AJ, Naidenko OV, Attinger A, Koning F, Lena CJ, Xiong Y, et al. T cell responses modulated through interaction between CD8alphaalpha and the nonclassical MHC class I molecule, TL. Science. 2001;294(5548):1936–9. PubMed PMID: 11729321.

41. Teitell M, Mescher MF, Olson CA, Littman DR, Kronenberg M. The thymus leukemia antigen binds human and mouse CD8. J Exp Med. 1991;174(5):1131–8. PubMed PMID: 1834760.

42. Weber GF, Ashkar S, Glimcher MJ, Cantor H. Receptor-ligand interaction between CD44 and osteopontin (Eta-1). Science. 1996;271(5248):509–12. Epub 1996/01/26. PubMed PMID: 8560266.

43. Bhagat G, Naiyer AJ, Shah JG, Harper J, Jabri B, Wang TC, et al. Small intestinal CD8+TCRgammadelta+NKG2A+ intraepithelial lymphocytes have attributes of regulatory cells in patients with celiac disease. J Clin Invest. 2008;118(1):281–93. Epub 2007/12/08. doi: 10.1172/JCI30989. PubMed PMID: 18064301; PubMed Central PMCID: PMCPMC2117760.

